# MPRAbase: A Massively Parallel Reporter Assay Database

**DOI:** 10.1101/2023.11.19.567742

**Authors:** Jingjing Zhao, Fotis A. Baltoumas, Maxwell A. Konnaris, Ioannis Mouratidis, Zhe Liu, Jasmine Sims, Vikram Agarwal, Georgios A. Pavlopoulos, Ilias Georgakopoulos-Soares, Nadav Ahituv

## Abstract

Massively parallel reporter assays (MPRAs) represent a set of high-throughput technologies that measure the functional effects of thousands of sequences/variants on gene regulatory activity. There are several different variations of MPRA technology and they are used for numerous applications, including regulatory element discovery, variant effect measurement, saturation mutagenesis, synthetic regulatory element generation or characterization of evolutionary gene regulatory differences. Despite their many designs and uses, there is no comprehensive database that incorporates the results of these experiments. To address this, we developed MPRAbase, a manually curated database that currently harbors 129 experiments, encompassing 17,718,677 elements tested across 35 cell types and 4 organisms. The MPRAbase web interface (http://www.mprabase.com) serves as a centralized user-friendly repository to download existing MPRA data for independent analysis and is designed with the ability to allow researchers to share their published data for rapid dissemination to the community.

## Introduction

Since the initial sequencing of the human genome, millions of *cis*-regulatory elements with putative roles in transcriptional gene regulation have been identified (Encode Project Consortium 2012; Thurman et al. 2012). Following up on their annotation, a major challenge has been to functionally characterize these elements. Massively parallel reporter assays (MPRAs) were built on the framework of the classic reporter assay. In this framework, the assayed sequence is placed in front of a reporter gene for promoter assays and also a minimal promoter for enhancer assays (**Fig. 1**). If the sequence itself has regulatory activity, it will turn on the reporter gene. To overcome the one-by-one testing limitation of these classic reporter assays, MPRAs add a DNA barcode that is transcribed if the sequence has regulatory activity and can be measured via RNA-sequencing (RNA-seq), providing a way to examine the functional effects of thousands of sequences in parallel (Patwardhan et al. 2009; Inoue and Ahituv 2015; Patwardhan et al. 2012; Melnikov et al. 2012; Agarwal et al. 2023) (**Fig. 1**). In recent years, the rapidly declining cost of DNA synthesis and sequencing have led to the growing popularity in the use of MPRA experiments and rapid accumulation of MPRA data.

**Figure 1.**
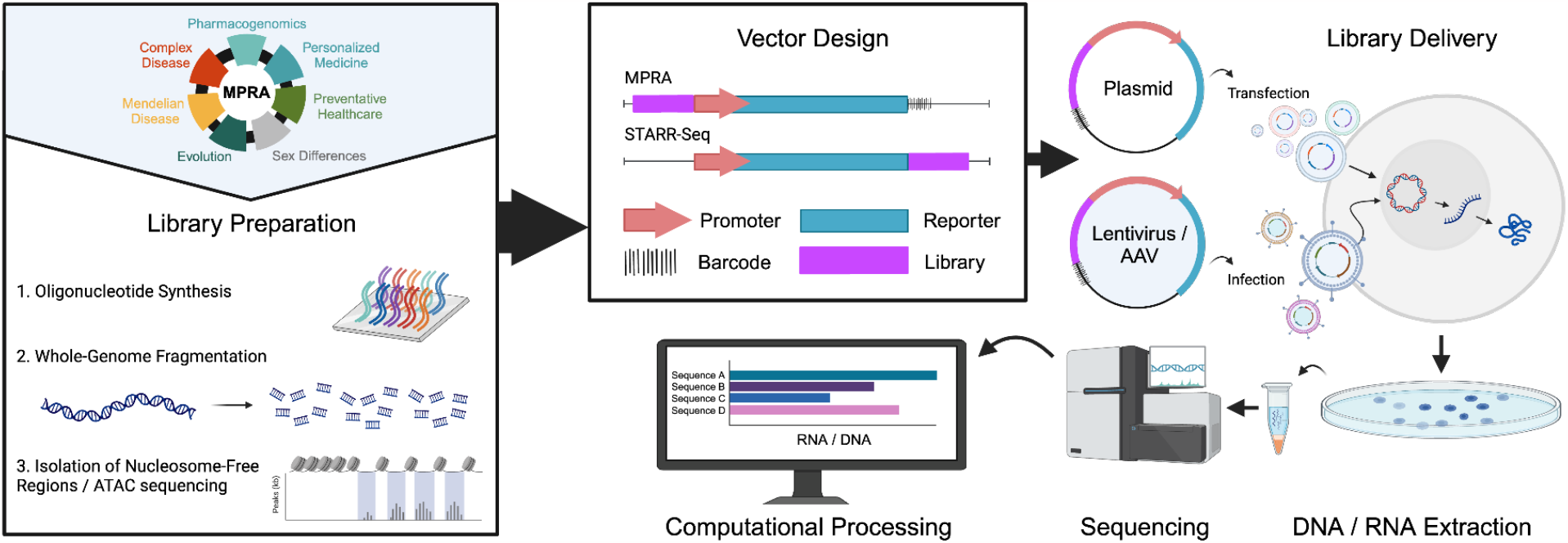
Schematic illustration of MPRA experiments. Various MPRA uses are described in the top left. MPRA libraries are usually prepared via oligonucleotide synthesis, genome fragmentation or isolation of nucleosome free or ATAC-seq regions. These sequences are then cloned into an MPRA vector along with a unique barcode, while in STARR-seq, the assayed sequence is used as the barcode. MPRA are then delivered to the cells either via transfection or viral infection. DNA and RNA are then extracted and the barcodes are sequences and activity scores are provided.

Since their invention more than a decade ago (Patwardhan et al. 2009), there has been a rapid emergence of novel variations of MPRA technology (**Fig. 1**) that differ by: 1) The positioning of the tested element and barcode relative to the reporter. For example, STARR-seq tests an element within a reporter’s 3′ UTR (Arnold et al. 2013); 2) Library generation. Libraries can be generated by the synthesis of pre-defined oligonucleotide sequences (Smith et al. 2013; Patwardhan et al. 2009, 2012; Melnikov et al. 2012; Agarwal et al. 2023), input of natural sequences from whole-genome fragmentation (van Arensbergen et al. 2016, 2019; Liu et al. 2017; Kvon et al. 2014; Arnold et al. 2013), isolation of nucleosome free regions (Murtha et al. 2014), or through the use of ATAC-seq (Wang et al. 2018); 3) The method of library delivery. MPRA libraries have been delivered to cells by transfection (Patwardhan et al. 2012; Melnikov et al. 2012; Kircher et al. 2019; Johnson et al. 2018; Liu et al. 2017), adeno-associated virus (AAV)-based MPRAs (Shen et al. 2016; Lambert et al. 2021; Chan et al. 2023) and lentivirus-based MPRAs (Inoue et al. 2017) that allow the integration of elements into the genome; 4) Computational processing tools, whereby the collected data are processed into activity scores, with appropriate processing pipelines chosen according to experimental design features (Lee et al. 2020; Gordon et al. 2020; Ashuach et al. 2019; Kim et al. 2021; Georgakopoulos-Soares et al. 2017). Experiments directly evaluating the aforementioned MPRA design choices have revealed a general consistency in measured element activities (Inoue et al. 2017; Klein et al. 2020).

Despite the exponential growth of published MPRA datasets, to date there is no centralized repository that aggregates the results of such data. To address this shortcoming, we introduce MPRAbase (http://www.mprabase.com), a database that harbors 129 experiments, encompassing a total of 17,718,677 sequences tested across 35 cell types and 4 organisms. In addition to storing published data, MPRAbase provides a platform that will make it easy for users to deposit new MPRA data and rapidly disseminate it to the functional genomics community.

MPRAbase has processed high throughput experiments across 50 studies. For each study, we provide the PMID and a link to the original publication. We also provide the mean expression score of each sequence, along with the expression score of each sequence for every replicate in the same format. MPRAbase offers an advanced search option, in which the user can search based on the coordinates of interest, the technique used (MPRA/STARR-seq and their variations), organism, cell type or motif. In addition, we provide an integration with the UCSC Genome Browser where available. MPRAbase also includes a filter for MPRAs carried out on synthetic sequences that do not exist in a specific genome. Finally, MPRAbase includes a separate tab for saturation mutagenesis MPRA that allows the selection of different regulatory elements, variants and variant scores.

## Results

### Database overview

Our goal with MPRAbase was to provide a central repository for all published MPRA and STARR-seq experiments. MPRAbase is designed to collect MPRA and STARR-seq data from different experiments, organisms and assays and provide them with a user-friendly web-based interface and to be able to easily download the data. The data provided in the database include the activity of sequences, measured as RNA/DNA ratio and provided separately for experimental replicates and associated correlation plots, metadata and statistics. MPRA experiments are divided into three types; i) standard MPRA, ii) synthetic MPRA, and iii) saturation MPRA experiments. Standard MPRAs are further subdivided into plasmid-based MPRAs, lentivirus-based MPRAs and STARR-seq.

### Collection of MPRA studies

For the development of our database, we scanned the literature using keywords and terms associated with MPRA experiments, resulting in the collection of 129 experiments. Studies were organized based on the organism of the assayed sequence, cell origin and type and experiment type. In total, MPRA experiments across 4 organisms, 35 cell types/tissues and 8 MPRA library types were downloaded, analyzed and presented in MPRAbase (**Fig. 2A-C**). The size of the MPRA libraries varies between 98 and 16,092,560 sequences, with 8,384 being the median number of sequences per experiment tested. The total number of DNA sequences available across all the MPRA experiments in MPRAbase is currently 17,718,677.

**Figure 2.**
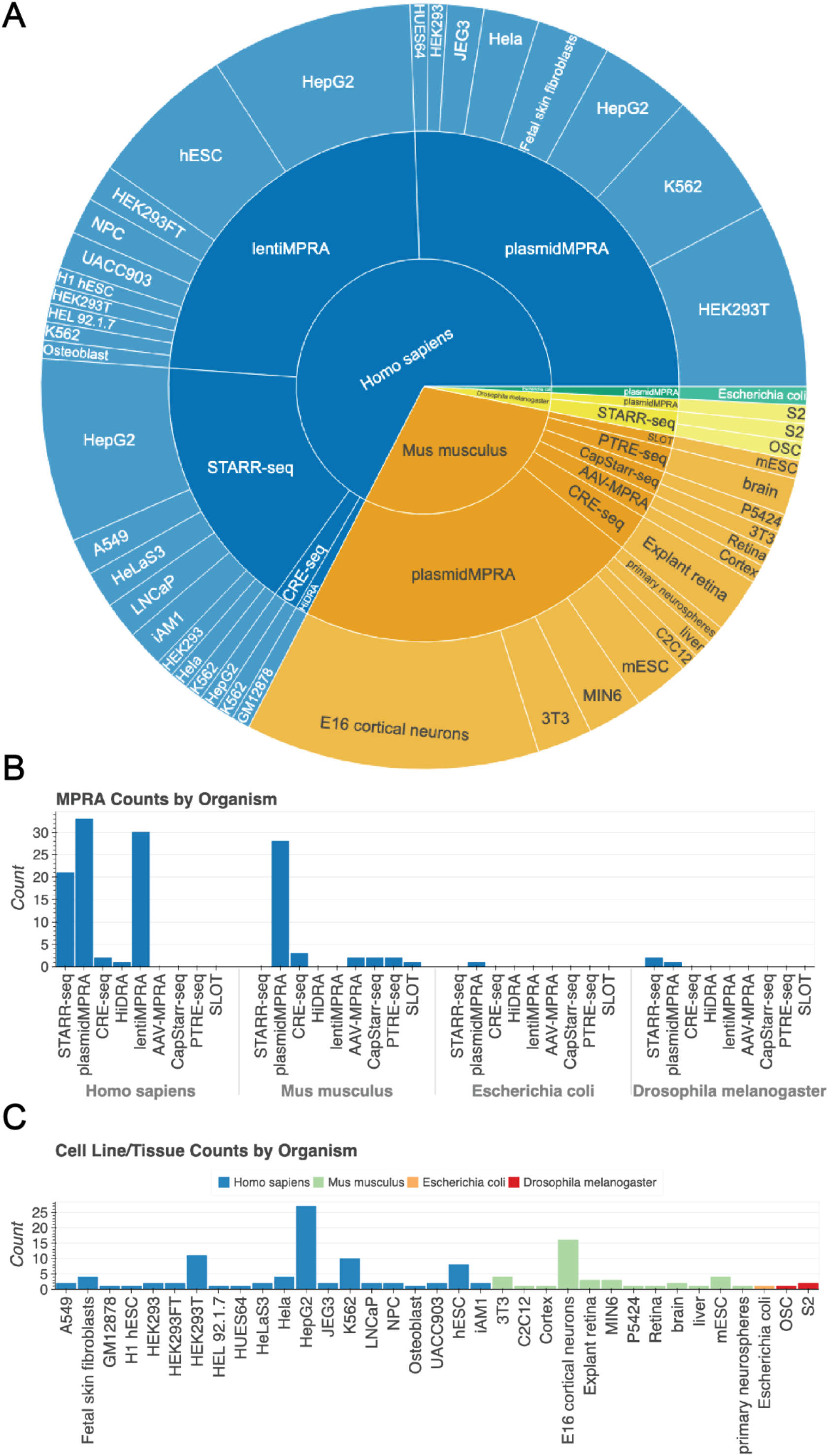
Visualization of MPRAbase summary statistics. *(A)* Pie chart displaying the break-down of MPRAbase by different species, MPRA type, and cell-type. (*B)* Bar plot showing the amount of experiments conducted for each species based on assay type. (*C)* Bar plot showing the amount of MPRA experiments carried out in each cell/tissue type for each species.

### Processing of MPRA data and expression quantification

In MPRA experiments, expression levels are generally quantified as the logarithmic DNA to RNA counts ratio with higher log-ratio reflecting increased *cis*-regulatory activity (Gordon et al. 2020). Data associated with each MPRA experiment across the collected studies were assembled and processed to provide the logarithmic ratio of RNA/DNA counts for each biological replicate and for the mean RNA/DNA expression levels across replicates.

MPRAbase also provides quality controls, including graphs for the correlation between biological replicates. The first type of plots compares the log RNA ratio between replicates, the second the log DNA ratio between replicates and the third the log RNA/DNA ratio correlations between biological replicates.

Library categories include plasmid-based MPRA experiments, lentivirus-based MPRA experiments (Gordon et al. 2020) and STARR-seq experiments (Muerdter et al. 2015). For lentiMPRA and plasmid-based MPRA experiments, the RNA/DNA ratio is provided for the elements of the library design, whereas for STARR-seq experiments genome-wide RNA/DNA ratios are quantified across retrieved coordinates.

### MPRAbase website and web-interface

The MPRAbase website contains interactive pie charts, tables and drop-down menus that enable the selection of MPRA experiments based on organism, cell type and library strategy used (**Fig. 3A-E**). The user can select multiple combinations of the aforementioned groups, for which an interactive table is presented with the individual samples. MPRAbase also provides a search bar to search regions by chromosome coordinates (**Fig. 3F**). Therefore, MPRAbase provides an additional functionality, in which the user can examine *cis*-regulatory activity for particular loci of interest. A set of coordinates for a reference genome of interest can be inserted by the user for which all MPRA sequences across experiments and cell types will be returned.

**Figure 3.**
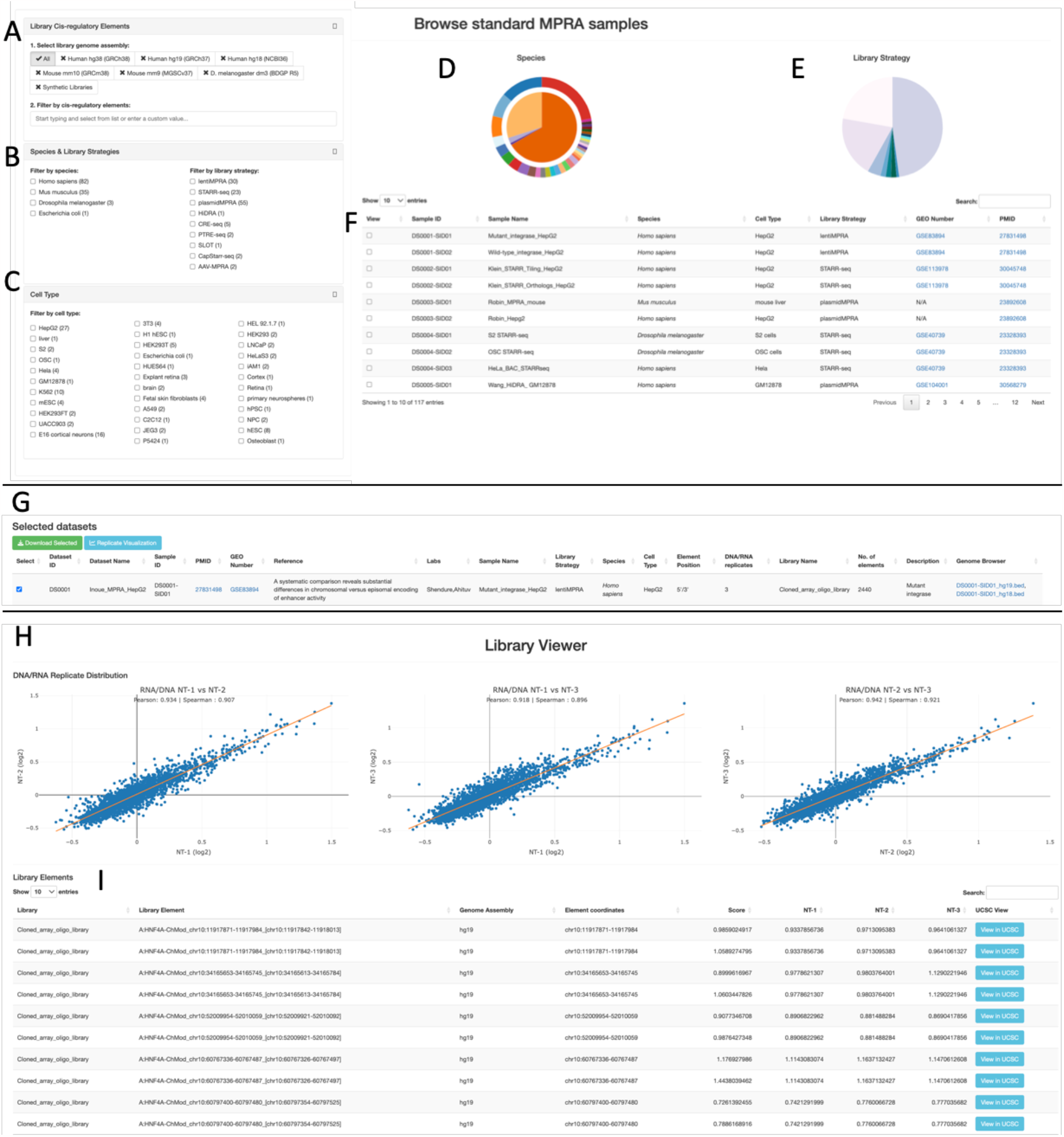
MPRA browser design. *(A)* Selection based on genome assembly or synthetic sequences with the option to search for specific loci of interest. (*B)* Selection based on species of interest or library type for different types of MPRAs to display by using a clickable box option. *(C)* Selection of cell-types of interest to display by using a clickable box option. *(D)* Pie-chart displaying MPRA dataset distribution by species. (*E)* Pie chart displaying MPRA category break-down to select from. (*F)* Table displaying the set of MPRA experiments in MPRAbase, with associated metadata including species, cell type, the library design used, the Gene Expression Omnibus (GEO) number of the experiment when available and the PubMed ID of the relevant publication. (*G)* Datasets selected can be visualized or the associated tables can be downloaded for further research. *(H)* Scatter-plots displaying pairwise correlations between replicates for RNA/DNA ratios. Line of best fit is displayed in yellow. Both Pearson and Spearman correlations are shown. *(I)* Table displaying *cis*-regulatory element activity for each tile, for the selected experiment. *Cis*-regulatory activity is displayed separately for each replicate in each column and a column of the combined activity is also displayed.

For each sample in the interactive table the sample ID, organism, cell type, library strategy, Gene Expression Omnibus (GEO) number and PubMed ID (PMID) of the experiment are provided. The identifiers associated with GEO and PMID entries have embedded clickable hyperlinks that can take the user to the associated studies and raw sequencing experiment databases. The selected data can be downloaded for further processing by the user. The format of the download is a zipped folder containing the sequence/RNA/DNA tables for the studies selected and a metadata file, which provides additional information for the selected studies.

MPRA is also carried out using synthetic sequences that may not exist in a certain genome. This is usually carried out in order to better understand the regulatory code, to design regulatory elements that can drive tissue/cell type specific expression, or sequences that respond to certain factors. To portray these in MPRAbase, we provide a specific filter called “Synthetic MPRA”.

### Saturation Mutagenesis

MPRAbase contains a separate tab for saturation mutagenesis MPRA (**Fig. 4**). In these MPRA experiments, a specific sequence is mutated and the effect of these numerous mutations is tested in parallel using MPRA. As it measures variant effects across the same sequence, we provide the ability to select the different promoters/enhancers that were tested using this approach by the name they were given in the experiment (**Fig. 4A-B**). Once the promoter/enhancer is selected, MPRAbase allows to select specific coordinates in the regulatory element for the variant scores, the number of unique tags, the log2 variant expression and p-value (**Fig. 4C-D**). Data is provided both as a table and also as a ‘lollipop figure’ that shows the different mutations and their effects (**Fig. 4C-D**).

**Figure 4.**
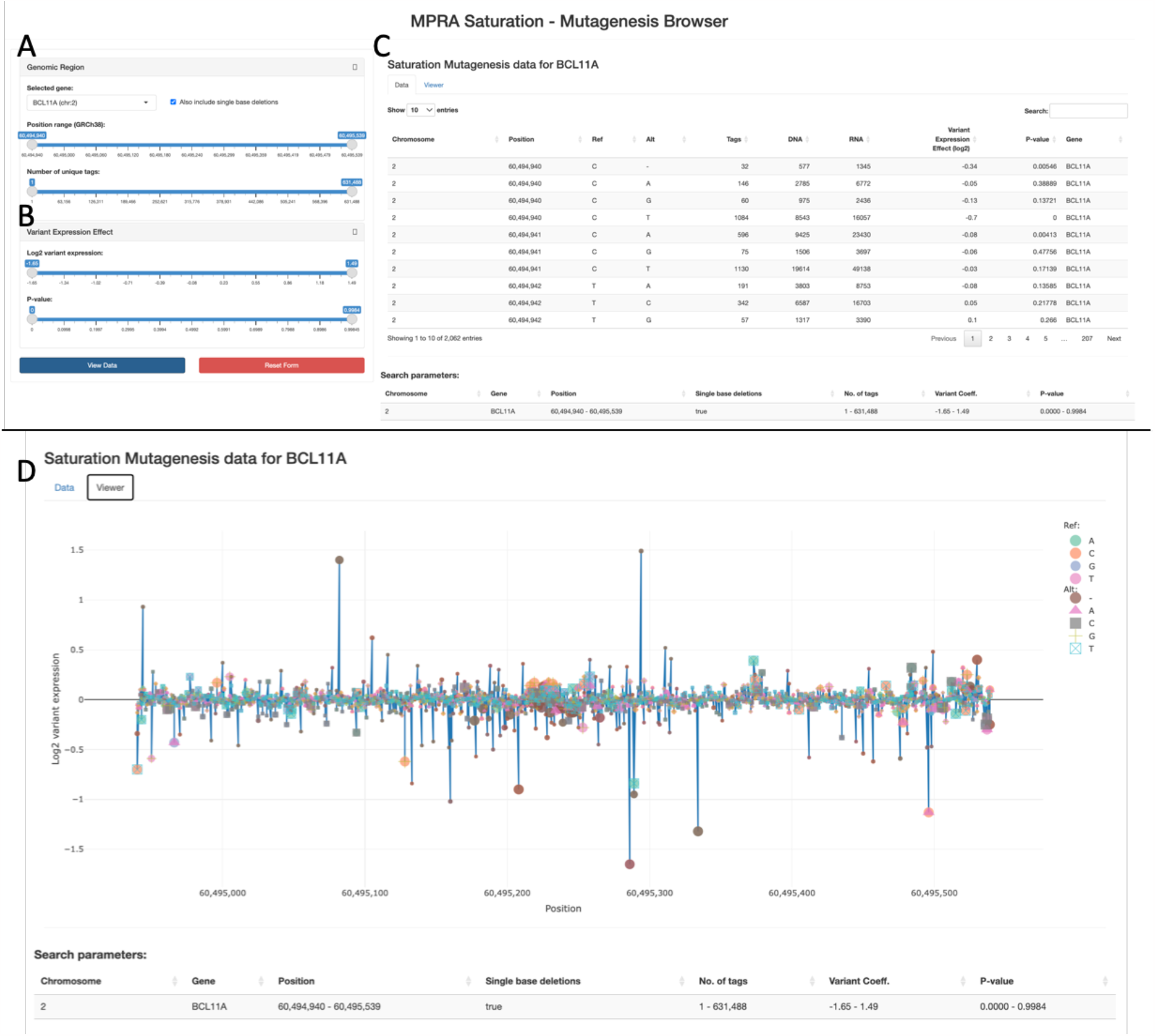
Saturation mutagenesis MPRAs on MPRAbase. *(A)* Drag-bar for selecting variants based on their position or number of unique barcodes associated with a variant. (*B)* Drag-bar for selecting based on variant expression effects. *(C)* Table displaying individual mutations for the selected locus and their effect on expression. (*D)* Lollipop graph displaying individual mutations for the selected locus and their effect on expression. Mutations are colored by Reference and Alteration type.

### Documentation and Help pages

The website has a “Documentation” page which provides information about MPRA experiments to introduce potential users to the MPRA technology and its utility. A help page is also provided, which provides explanations for the different functionalities of the database.

## Discussion

The recent advances in DNA synthesis and sequencing costs have enabled the widespread adoption of MPRA technologies. MPRAs and other related assays can provide insights into the roles of disease-associated variants, can be used to gain insights in *cis*-regulation and have been implemented in studies of primate evolution, while they can also be used to examine synthetic sequences, with potential applications as therapeutic molecules (Whalen et al. 2023; Georgakopoulos-Soares et al. 2023; Deng et al. 2023; Georgakopoulos-Soares et al. 2022; Arnold et al. 2013; Agarwal et al. 2023). Here, we generated MPRAbase that allows the user to view and analyze MPRA datasets in one location.

MPRAbase provides a curated database for MPRAs, consisting, at launch, of 129 experiments, for 36 cell types, across 4 organisms. The website is user-friendly and interactive, enabling users to select studies based on a list of criteria, including organism, cell type and MPRA library type. For each study, we provide quality control metrics, enabling users to decide if the selected studies meet their quality requirements. We plan to have MPRAbase updated regularly to accommodate the increasing number of available MPRA experiments. With a continuously updated, comprehensive characterization of MPRA experiments across organisms and cell types, we believe MPRAbase will be a valuable resource to better understand gene regulatory grammar, illuminate the consequences of non-coding mutations and be used to gain insights into evolutionary facets. We therefore anticipate this resource will have a broader impact on our broader understanding of genetics.

## Author Contributions

J.Z., V.A., I.G.S., and N.A. conceived of the study; J.Z.,. F.A.B.,. M.A.K,. V.A., I.M., Z.L., J.S., and I.G.S performed the computational analyses; J.Z.,. F.A.B.,. M.A.K,. and I.G.S. generated the visualizations; G.A.P., I.G.S., and N.A. provided resources; I.G.S., and N.A. wrote the manuscript with input from all authors; V.A., G.A.P., I.G.S., and N.A. supervised the study.

## Methods

### MPRA experiments in database

Publications with MPRA, STARR-seq or other related assays were systematically collected. MPRA data were retrieved and manually curated. Curation included the collection of the PMID, organism name, cell type, experiment type and library strategy used. Whenever available the GEO ID was also integrated. Correlation analyses were performed between replicates for each study for RNA/DNA ratios. For human MPRAs, MPRA coordinates in hg18 or hg19 were revised to hg38 using the UCSC liftOver tool (Kuhn et al. 2007); in mice mm9 coordinates were converted to mm10 .

### Database implementation

MPRAbase contents are organized in a relational SQLite database (https://www.sqlite.org/). The user interface was implemented using R (https://www.r-project.org/) and the R/Shiny framework. Server-side operations are mainly handled by R. Data visualization and graphs are generated using the R/DT and R/plotly packages (Sievert 2020). MPRAbase is available online without fees for academic usage. The database is updated in regular 3-month intervals, as new MPRA studies become available.

